# Automated Rational Strain Construction Based on High-Throughput Conjugation

**DOI:** 10.1101/2020.11.24.396200

**Authors:** Niklas Tenhaef, Robert Stella, Julia Frunzke, Stephan Noack

**Affiliations:** Institute of Bio- and Geosciences – IBG-1: Biotechnology, Forschungszentrum Jülich GmbH, 52425, Jülich, Germany; Bioeconomy Science Center (BioSC), Forschungszentrum Jülich, Jülich, 52425, Germany

**Keywords:** Laboratory automation, Strain construction, Molecular cloning, Transcription factor-based biosensors, DNA assembly, High-throughput conjugation

## Abstract

Molecular cloning is the core of Synthetic Biology, as it comprises the assembly of DNA and its expression in target hosts. At present, however, cloning is most often a manual, time-consuming and repetitive process that highly benefits from automation. The automation of a complete rational cloning procedure, *i.e.*, from DNA part creation to expression in the target host, involves the integration of different operations and machines. Examples of such workflows are sparse, especially when the design is rational (*i.e*., the DNA sequence design is fixed, and not based on randomized libraries) and the target host is less genetically tractable (*e.g.*, not sensitive to heat-shock transformation). In this study, an automated workflow for the rational construction of plasmids and their subsequent conjugative transfer into the biotechnological platform organism *Corynebacterium glutamicum* is presented. The whole workflow is accompanied by a custom-made software tool. As an application example, a rationally designed library of transcription factor biosensors based on the regulator Lrp was constructed and characterized. A sensor with an improved dynamic range was obtained, and insights from the screening provided evidence for a dual regulator function of *C. glutamicum* Lrp.

## Introduction

Microbial production of bulk and fine chemicals is a vital part of a more sustainable global economy. To foster this development, both the fundamental understanding of microbial life and its engineering to fulfill society’s needs must be advanced. In recent years, the employment of a Design-Build-Test-Learn (DBTL) cycle has been proposed as a tool to achieve this [1]. Molecular cloning plays an important role in this cycle since it allows for the generation of new genotypes with different properties to explore.

Multiple software tools have been developed to aid the *in-silico* cloning of genotypes, in numbers that easily exceed the amount that could be manually made in the laboratory [2]. It is now feasible to design many hundreds of genotypes in a short amount of time. For example, a production pathway of five genes each with three different ribosomal binding sites (RBSs) already results in 3^5^ = 243 variants. However, the relatively easy design of such a project now shifts the bottleneck to the actual *in vitro* creation of these sequences, and their expression in the desired industrial host [3].

This problem can be addressed by using one-pot assembly & screening approaches, *i.e.*, to obtain many but not necessarily all variants and use a screening assay to select for the best performers [4]. However, these approaches have one drawback: Knowledge is only obtained from the few variants that were successfully constructed and isolated from the one-pot reaction. More difficult to construct variants are less abundant in such a reaction and are therefore less likely to be screened for their properties. For many fundamental biological questions, this is not an ideal solution. Here, rational strain construction with complete traceability of all varieties within all steps of the workflow is necessary, as is realized by classical molecular cloning.

At present, however, molecular cloning is most often a manual, time-consuming and repetitive process that would highly benefit from automation [5]. The automation of a rational cloning workflow has multiple benefits. The hands-on time of the experimenter spend on strain construction can be drastically reduced. Together with the higher throughput that can be realized, this largely increases the quantity of constructs that can be produced in time. Furthermore, automation introduces standardization of the process, by removing random variations from the process. The processes can also be more easily monitored and analyzed, making it easier to find room for improvement. Thus, automation can increase both the quantity and quality of cloning workflows.

In recent years, microfluidics has been used to automate cloning processes [6, 7]. Here, tailor-made microfluidic chips are used to provide liquid separation for and liquid transfer in between the single unit operations. While this technique comes with the advantage of combining many unit operations in one device and the ability to scale up to a high number of experiments per chip, highly specialized infrastructure and personnel is needed to fabricate the chips and to conduct the experiments.

A more modular and accessible solution is the employment of standard liquid handling systems and, if needed, auxiliary devices. Such a system can be rather complex and capable of performing a high number of experiments with varying tasks [8–10], but also more cost-efficient solutions are available [11]. A recently presented strategy uses the natural transformation capacity of specific bacteria, resulting in a highly efficient and easy to use cloning method [12]. However, this approach is limited to the few organisms which actively take up exogenous DNA.

Most automated cloning workflows published so far focused on working with *Escherichia coli* [13]. *E. coli* is a well-established host for molecular cloning that is very genetically tractable. Many genetic tools have been developed and optimized for use in *E. coli* and comprehensive omics data is available. However, *E. coli* is not always an ideal host for industrial processes, due to its low stress tolerance and risk of phage infection [14]. Therefore, it would be beneficial to extend automation platforms to include the engineering of other microorganisms that are less genetically tractable.

*Corynebacterium glutamicum* is a widely used industrial bacteria [15] that is more difficult to engineer than *E. coli*, due to its resistance to automation friendly transformation processes such as heat shock transformation. While some steps have been taken in this direction [16], they usually fall short in automating the process of transforming the actual target organism, most often because electroporation, which is the method of choice for such organisms, is not easily accessible for automation.

This study presents a complete workflow for automated rational strain construction of heat shock resistant microorganism. All work was carried out using standard liquid handling systems. PCR and Gibson assembly were used to construct a library of 96 plasmids. Automated protocols were developed for heat-shock transformation of *E. coli* as a shuttle system. Colony PCR and sequencing were used as quality control. For the final step of transforming *C. glutamicum*, a novel high-throughput conjugation workflow was d eveloped. Conjugation is an well-described and highly relevant technique for transformation of bacteria [17]. High efficiency can be obtained, but the method is usually laborious because of the use of agar plates and filter papers. For this study, the operation was simplified and made accessible to automation by using centrifugation. The whole workflow was accompanied by a custom-made software tool to keep track of all constructs and their status within the process.

As an application example, the assembly and expression of different Lrp biosensor variants in *C. glutamicum* is shown. The Lrp biosensor was previously developed for the detection of L-methionine and branched-chain amino acids in *C. glutamicum* [18]. This sensor couples intracellular L-methionine and branched-chain amino acid concentration to expression of *eyfp*, encoding a fluorescent reporter protein. Increased intracellular concentration results in a higher fluorescent signal. In general, biosensors are relatively modular in their design, their characteristics can be changed by modifying, for example, ribosomal binding sites and promoter length [19, 20]. Furthermore, by design they provide a direct and easily measurable relationship between genotype and phenotype, *i.e.*, a change in Lrp sensor design will likely result in a different, measurable, fluorescence output. Therefore, the rational design of different Lrp biosensors was chosen as an application example demonstrating the strength of our cloning automation approach.

## Results and Discussion

### Workflow for automated genetic engineering

The automated cloning workflow developed in this study (Figure 1) can be divided into two stages: Plasmid assembly and amplification in *E. coli*, and transfer of the plasmid to the target organism, in this case *C. glutamicum*. Plasmids were constructed by designing the parts *in silico*, building fragments by PCR and integration into a backbone vector by Gibson assembly. Gibson assembly was chosen because it allows for scarless assembly of plasmids [21]. The subsequent transformation into *E. coli* was done by heat-shock. Resulting clones were stored by cryo-conservation and a first quality control step was done by colony PCR. This method was chosen because of its cost-efficiency and short run times. The results from this colony PCR influenced the next steps: Only *E. coli* clones with a positive colony PCR result, *i.e.*, with a resulting DNA fragment of the expected size, indicating the successful assembly of a fragment into the backbone, were considered for automated plasmid preparation and, in parallel, conjugation into *C. glutamicum*. Purified plasmids were used for sequencing as a final quality control. Afterwards the automatically constructed strains were screened to characterize the altered properties. Most importantly, each unit operation was designed to be self-contained, so it can be used detached from the workflow.

**Fig. 1.**
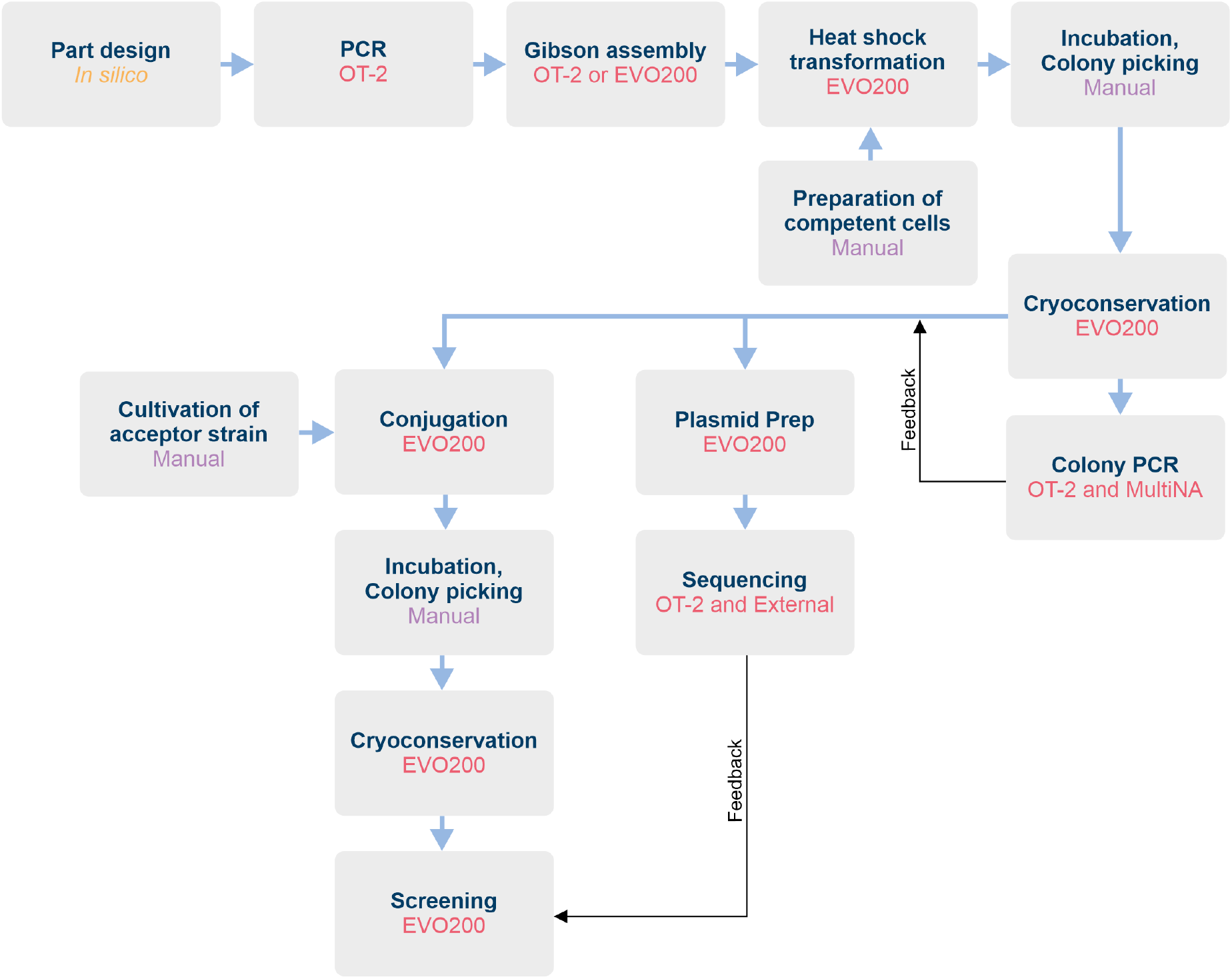
Overview of the automated genetic engineering workflow. Each box describes one unit operation and informs about the device or entity used for performing this operation. OT-2: Opentrons OT-2 liquid handling system. EVO200: Tecan EVO200 liquid handling system. MultiNA: Shimadzu MCE-202 MultiNA chip electrophoresis system.

Automated unit operations were carried out or supported by a low-cost liquid handling device, the Opentrons OT-2, and a more sophisticated liquid handling platform based on a Tecan EVO 200 described earlier [22, 23]. Both systems are fundamentally different: The OT-2 uses a setup of two air displacement pipettes working with disposable tips. Available pipettes have a volume range similar to a manual pipette and the operator has to select the ones suitable for the experiment. It does not have a robotic manipulator; therefore, it is not able to change the location of plates, *e.g.*, to place a plate on a heating block or in a centrifuge. The deck can hold up two nine MTPs. The model used for this study was not equipped with an option to cool labware. A custom-made cooling rack which could be filled with ice and fitting adapters were designed and 3D printed (see Supporting Information) to cool down reagents when necessary (*e.g.*, for Gibson assembly). The EVO 200, in the configuration used in this study, has a liquid-based liquid handling system, spanning a volume range of 3 to 990 *µ*L. It is equipped with a centrifuge, a cooling carrier, and a heater/shaker-unit, both accessible for the integrated robotic manipulator arm. The deck can hold 15 or more MTPs, depending on the carriers used. For spotting of bacterial cultures onto round 100 mm petri dishes, a custom-made adapter was designed and 3D printed (see Supporting Information).

The decision to use two different liquid handling systems was motivated by the need to use lab equipment most efficiently: Whenever possible, the OT-2 was used to carry out a unit operation. This saved time on the more expensive EVO 200, which thus could be used for more demanding experiments. The unit operation “colony picking” was done manually in this study. It can be automated by dedicated devices, but none of those were available at the time.

### Rational combinatorial design of transcription factor-based biosensors

To demonstrate the applicability of our workflow, different versions of the Lrp biosensor were designed, constructed and expressed in *C. glutamicum*. The Lrp biosensor couples intra-cellular L-methionine, L-leucine, L-isoleucine and L-valine concentration to expression of the fluorescent reporter protein eYFP. To construct different versions of the Lrp biosensor, different parts of the sensor were modified. The Lrp biosensor consists of the *lrp* gene, which encodes the Lrp transcription factor, the divergently expressed *eyfp* gene, encoding the fluorescent r eporter, a nd t he i ntergenic r egion, w hich contains the *lrp* promoter and the *brnF* promoter upstream of *eyfp* and the *eyfp* RBS [18] (Figure 2). Different variants were designed for the *lrp* start codon and RBS, for the *brnF* promoter and for the *eyfp* RBS.

**Fig. 2.**
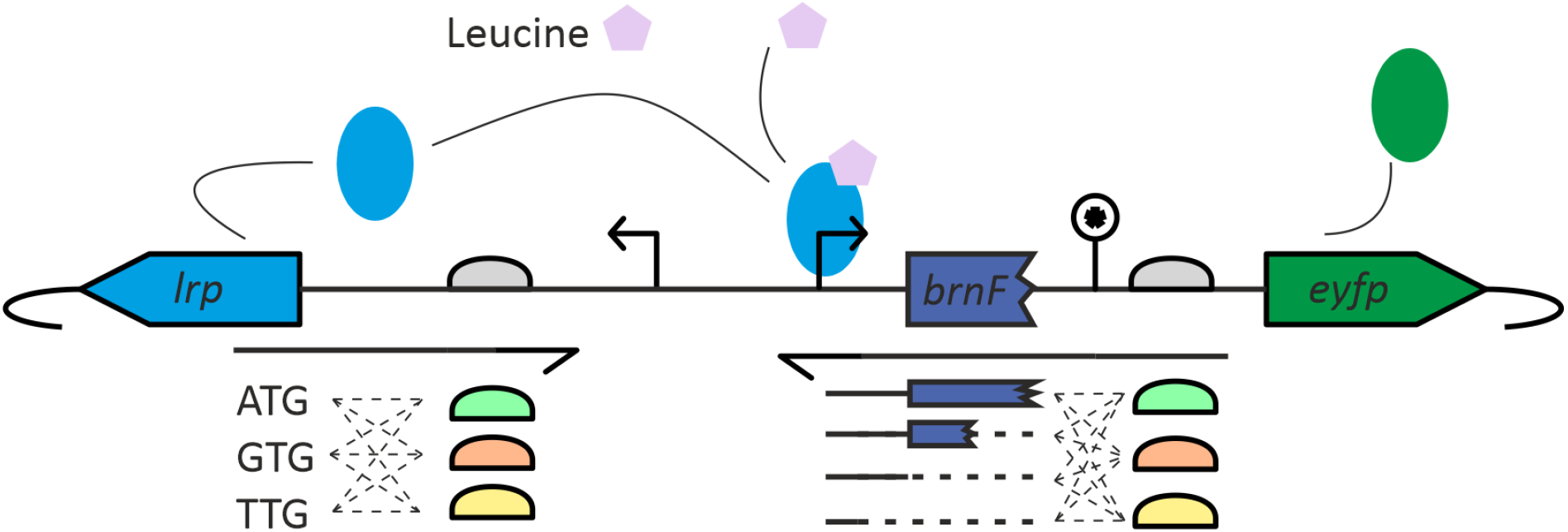
Graphical overview of the different Lrp biosensor variants that were designed for construction and expression in *C. glutamicum*. On the left side, 9 different primers were designed, containing combinations of different *lrp* start codons and *lrp* RBSs. On the right side, 12 different primers were designed, containing combinations of different *brnF* promoter sequences and *eyfp* RBSs.

For the *lrp* start codon, three variants were chosen, the native start codon ATG, which should result in the highest expression, the start codon GTG, which should result in lower expression, and TGT, which should result in very low or no expression. Each of these start codon variants were combined with three different *lrp* RBSs, again ranging from highest expression to low expression: GCTAAAATGG (strongest) GC-TATTGTGC (native, less strong) and CAATCCTACC (weakest). The combination of three start codons and three RBSs at the *lrp* side resulted in 3 × 3 = 9 different primers.

The *eyfp* gene was expressed under control of the *brnF* promoter, which contains binding sites for the Lrp protein [24]. Thus, by shortening or elongating the promoter sequence, the binding site(s) can be included or removed. Because *brnF* is transcribed as a leaderless transcript and the first part of *brnF* probably has a regulatory function [25]. Therefore, the *eyfp* promoter of the standard Lrp sensor includes the first 30 bp of *brnF*. In this study, four different variants were designed, starting at +30 (standard Lrp biosensor promoter) and shortened versions in steps of 15, *i.e.*, +15, 0 and −15. For the RBS of *eyfp* three variants were chosen, AAAAGGAGAT (strongest), AGAAGGAGAT (native, less strong) and ATCCGACCAT (weakest). The combination of four promoter lengths and three RBSs at the *eyfp* side resulted in 4 × 3 = 12 different primers. This design strategy would include 9 × 12 = 108 different variants of the Lrp biosensor in total.

Each variant can be constructed by a single PCR reaction, and all PCR reactions can use the same PCR template, plasmid pJC1-*lrp*-*brnF’*-*eyfp*. To assemble the biosensor plasmids, each PCR reaction is followed by a two-fragment Gibson assembly, in which the PCR product is ligated into a pEC-Tmob18-*lrp*-*eyfp* backbone, all PCR products were designed as such that they can be ligated into the same backbone.

### DNA assembly and transformation of *E. coli*

Before construction of the Lrp biosensor variants, a backbone plasmid was constructed by inserting *lrp* and *eyfp* into the mobilizable pEC-Tmob18 backbone. A short spacer that contained two NcoI restriction sites was inserted between the two genes. This sequence allowed for plasmid linearization, and insertion of different versions of the sequence variants containing different *lrp* start codons, *lrp* RBSs, *brnF* promoter lengths and *eyfp* RBSs, in the next step.

Out of the 108 variants, 96 were selected as an application example for the automated cloning workflow, because most automation devices are designed for 96 well multititer plates. Variant construction by PCR, Gibson assembly and *E. coli* heat shock transformation were performed directly after each other. The Opentrons OT-2 enabled automated pipetting of 96 PCR reactions in 85 minutes, after which the plate was manually transferred to a thermocycler. After thermocycling, the OT-2 was used to mix the 96 Gibson assembly reactions, which took 50 minutes. Incubation was done by manually transferring the plate to a thermocycler. Appropriate cooling of the Gibson master mix before the 50 °C incubation step turned out to be crucial, since otherwise no *E. coli* transformants were obtained. In our workflow, cooling was achieved by placing all labware in small 3D-printed boxes filled with ice. *E. coli* heat shock transformation was performed on a TECAN EVO200 liquid handling robot, which enabled a completely hands-off process, starting with the Gibson assembly mixtures and competent cells, and ending with the cells spotted on agar plates ready for overnight incubation. This process took 170 minutes. In total, PCR, Gibson assembly and transformation of 96 constructs was achieved in 8 hours with only 40 minutes hands on time.

The best way to implement the heat shock step was to pre-heat a V-bottom 96-well plate on a heating device, transfer 8 cell-plasmid mixtures at a time from the cooling carrier to the heated plate, incubate for 30 seconds, and transfer back to the cooling carrier. While transferring a complete plate from the cooling carrier to the heating device is faster, this gives less reliable heat transfer results. Plating out was done by concentrating the cell suspension by centrifugation, resuspending in LB medium and spotting on agar plates, *i.e.*, by pipetting 6 spots of 5 *µ*L on the same row of a single agar plate.

Additionally, parameters for heat shock transformation were tested using *E. coli* competent cells (NEB® 5-alpha Competent *E. coli*) using a pUC19 standard vector. Heat shock temperatures (37 °C, 42 °C, and 47 °C) and heat shock durations (0 to 30 seconds, in steps of 5 seconds) were tested. Surprisingly, little difference in colony forming units was observed between the conditions, with successful transformations for each condition. Eventually, conditions most similar to standard manual routines were chosen.

Transformed *E. coli* clones were counted after ~ 16 hours of plate incubation at 37 °C. A total amount of 273 clones were obtained, with at maximum of 12 clones per construct. The number of unique constructs for which a single clone was obtained was 60 out of 96. Colony PCR was done on at most 4 clones of each construct, to test if the PCR product was correctly inserted into the pEC-Tmob18-*lrp*-*eyfp* backbone in the Gibson assembly step. A total of 184 clones was tested, and results were analysed on a MultiNA, which allowed for easy and high throughput analysis of PCR band sizes for all constructs. From 184 clones, 172 had a PCR band with a length within 10 base pairs from the target length. If only unique constructs are considered, 55 out of 60 constructs were correct. To validate the results of the MultiNA, a classical agarose gel-electrophoresis was done for 16 colony-PCR products and the result compared to the MultiNA results (Figure 3). The obtained bands are highly comparable between the two methods, validating the use of the MultiNA as a replacement for classical gel-electrophoresis.

**Fig. 3.**
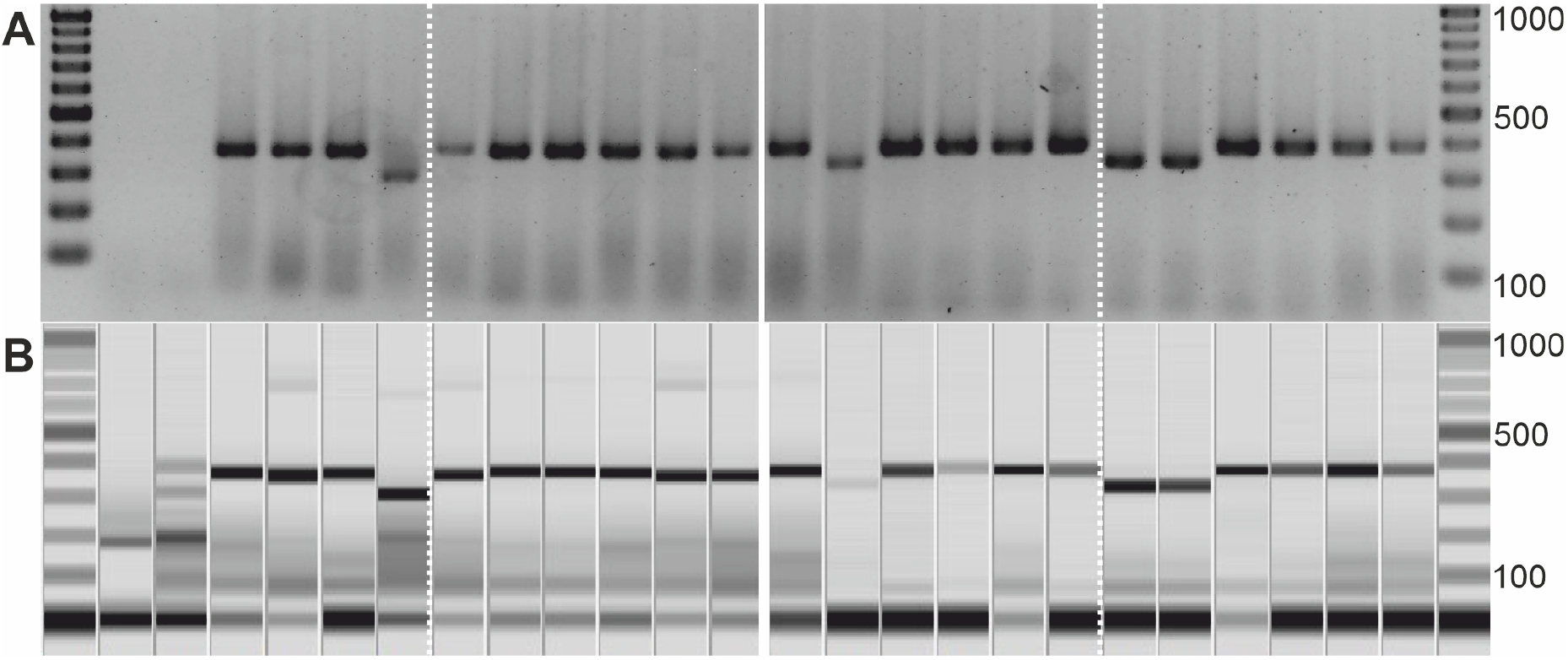
Comparison of standard gel electrophoresis and MultiNA capillary gel electrophoresis for Colony PCR analysis.(A) Annotated gel image. (B) Virtual gel image created from gel electrophoresis data. Linear DNA fragment sizes are shown on the right size, in basepairs (bp). NEB 1 kb Plus DNA Ladder was used as marker.

### Transformation of *C. glutamicum* and strain validation

Automating the transfer of plasmid into *C. glutamicum* provided a challenge, because no chemical protocols (*e.g.*, heat-shock) have been shown to work, and available high-throughput electroporation devices are not automation friendly. Conjugation is an alternative plasmid transfer method, in which a donor cell (*e.g.*, *E. coli*) transfers a plasmid into an acceptor cell (*e.g.*, *C. glutamicum*). For conjugation, cells need to be in close contact to each other, which is typically achieved by mixing the donor and acceptor cultures and spreading the mixture on an agar plate. This is usually on top of a filter, to allow for easy removal of the cells. Automated conjugation was implemented on the Tecan EVO200 by mixing the donor and acceptor cultures and performing a centrifugation step to avoid the plating out step, but to keep the cultures in close contact.

Initially, three different types of plasmid were considered for conjugation, pEC-Tmob18, pEC-Cmob18 and pEC-Tmob18 [26, 27], harboring a resistance marker for tetracycline, chloramphenicol and streptomycin, respectively. Consistent successful conjugation results were only obtained for pEC-Tmob18, which was therefore used for all experiments. Two parameters were investigated to optimize the conjugation protocol. First, the effect of cell concentration was tested, by combining different *C. glutamicum* starting culture optical densities (OD_600_) with different *E. coli* starting culture OD_600_ (Figure 4). The effect of the *C. glutamicum* starting culture OD_600_ was much larger than that of the *E. coli* starting culture OD_600_, with higher densities resulting in more clones. Furthermore, when using overnight cultures to conjugate the plasmids of 96 *E. coli* strains harboring different Lrp biosensor-plasmids, it was found that older overnight cultures (more than 20 hours) performed worse (47 clones, 38 unique) than fresh high density cultures (233 clones, 84 unique). This effect was likely due to the starvation effects of prolonged stationary phase. Finally, the effect of *C. glutamicum* growth media was investigated. Changing to BHI media instead of LB media for *C. glutamicum* preculture growth gave the best results, (995 clones, 96 unique, at least 4 clones per conjugation reaction).

**Fig. 4.**
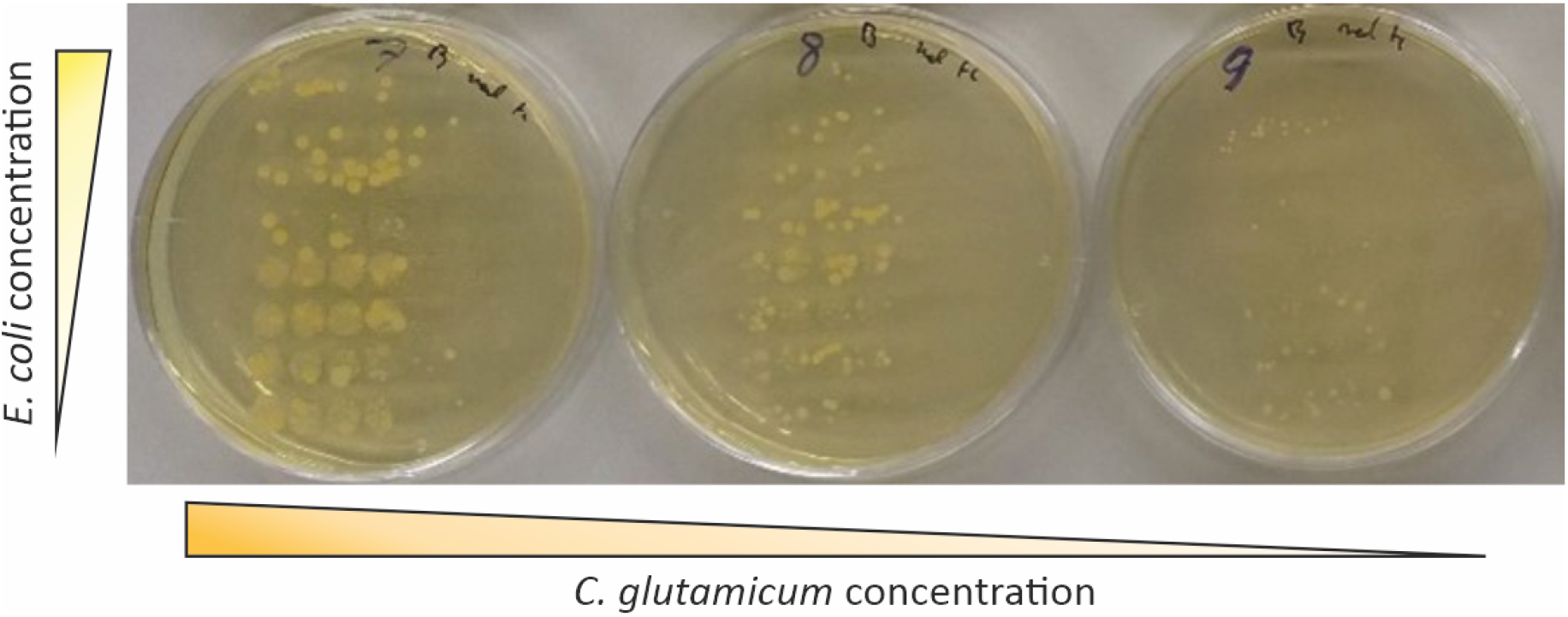
Effect of *E. coli* (donor) and *C. glutamicum* (acceptor) starting culture optical density on conjugation results. Each row on a single BHI agar plate represents multiple cultures from the same conditions. Less clones are visible with decreasing *C. glutamicum* starting culture optical densities, while the effect of *E. coli* optical density does not have a large effect.

Using the Tecan EVO200 platform, 96 conjugation reactions could be performed in 70 minutes, with less than 10 minutes hands on time. To transfer the biosensor variants to *C. glutamicum*, 96 *E. coli* cultures harboring 55 unique constructs were used for the conjugation protocol. At least 4 *C. glutamicum* clones per construct were obtained (995 clones in total). However, not all clones could grow in liquid cultures after being picked from the agar plate – an effect which is probably backbone-dependent. Picking 4 clones per construct into liquid BHI media resulted in 64 strains with one or more clones that could grow in liquid media, including 45 unique strains. Interestingly, at least some of the clones that were not able to grow in liquid BHI media did contain a plasmid, as confirmed with colony PCR. The reason for this unexpected growth behavior is not clear and should be investigated in the future.

### Plasmid preparation and validation

After conjugation, 45 unique constructs were transferred to *C. glutamicum*. To validate the sequence of these sensors-variants, the relevant plasmids were sequenced (externally) by sanger sequencing. Sanger sequencing required purified plasmids. Using the TACO software (as described below), the location of the *E. coli* clones harboring the plasmids that were successfully transferred to *C. glutamicum* was retrieved, and these clones were picked for overnight growth and plasmid purification. Using the Tecan platform, 45 plasmid purifications could be performed in 105 m inutes. Measuring 8 samples gave concentrations between 25 and 50 ng *µ*L^−1^ (elution volume 100 *µ*L), which is on par with plasmid amounts obtained with manual miniprep kits.

Sanger sequencing revealed that 36 out of the 45 plasmids had the expected sequence. From the 9 plasmids without a correct sequence, 4 were constructed using the same primer, and all missed a large part of the insert-sequence. This could indicate a problem with this primer, but also correct constructs were obtained using the same primer. For the other mutations, no correlation was found. However, a result of ~ 80 % correct sequences is not below results obtained using manual cloning procedures.

### TACO

All molecular cloning workflows r equire a c ertain e ffort in documentation of design ideas, results and storage locations of DNA fragments, plasmids, clones etc. With increased throughput, this task becomes increasingly difficult and cannot be handled anymore by standard techniques such as manually maintained (electronic) lab notebooks. To solve this problem, a Tool for Automated Cloning (TACO) was developed. Using Python 3.8 and building on top of the widely used libraries Pandas and Numpy, TACO provides the user with templates to keep track of different constructs, number of clones resulting from transformation experiments, and plate storage locations. All templates are provided as Microsoft Excel files for easy access and transfer to lab locations (*e.g.*, by printing). Submodules allow for expansion of TACO. One submodule was developed to analyze data from the capillary gel electrophoresis device MultiNA, used for colony PCR in this workflow. Another submodule was made to assist with the conjugation part of the workflow. To provide visual aids for the user, Graphviz was used to create directed acyclic graphs (DAGs) from both plasmid assembly and conjugation stages of the workflow. An illustration of such graphs can be found in Figure 5.

**Fig. 5.**
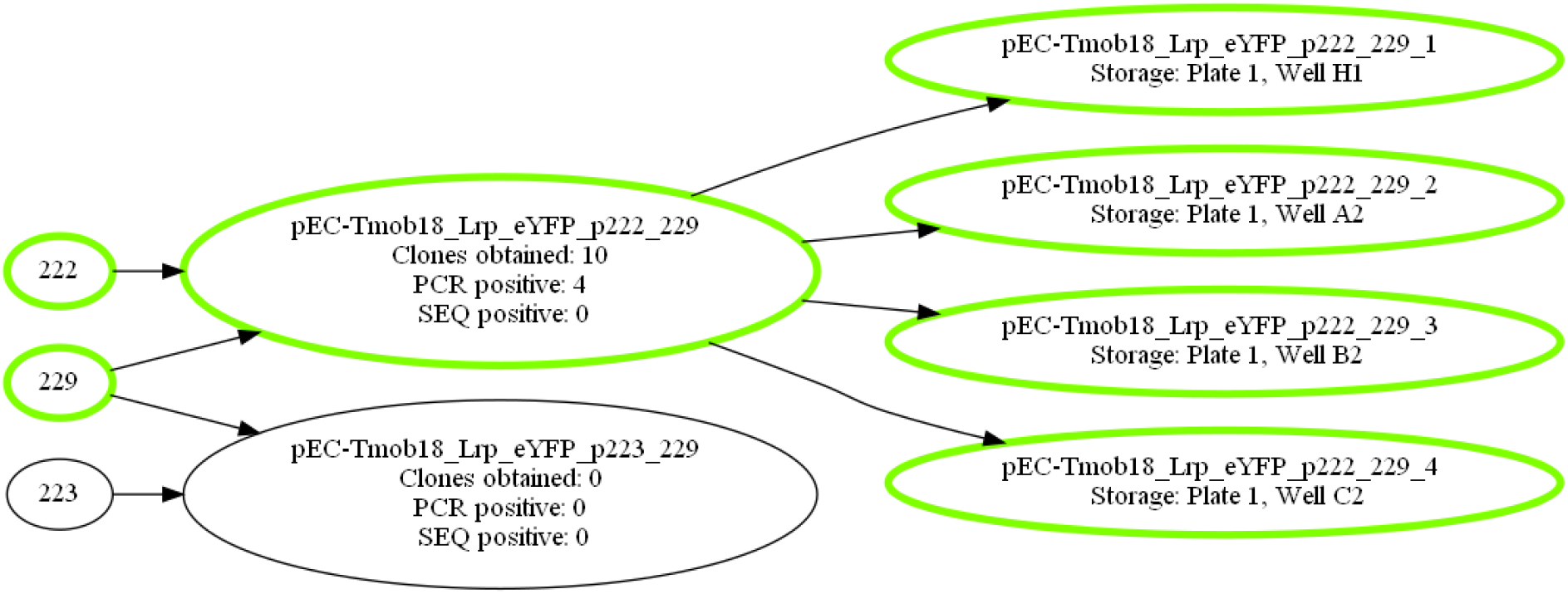
Part of the DAG for conjugation stage of the application example. Initial fragments, constructs, conjugation events and stored clones form nodes. Edges visualize their relationship. The user can choose to highlight successfully stored clones and their linage, as done here.

### Screening of novel biosensor strains

After the complete cloning workflow, 36 *C. glutamicum* strains with different versions of the Lrp sensor were obtained. During initial testing experiments, 8 more *C. glutamicum* strains with different, validated, Lrp sensors were obtained, bringing the total to 44. The sensor response was measured by growing the strains in microtiter plates, in presence or absence of 3 mM inducer (alanine-leucine dipeptide). Inoculation of main cultures and automated measurement was carried out on the Tecan EVO200 platform. Fluorescence output was measured, and fold-change was calculated (Figure 6). The highest fold-change was observed for the sensors with the strongest *lrp* RBS and start codon, *eyfp* RBS, and the longest promoter containing the first 30 bp of *brnF*. In general, the fold-change was highest when using a strong *lrp* RBS and start codon in combination with strong *eyfp* RBS. When the strength of either of the four elements was lowered, a decrease in fold-change could be observed. In contrast, when the *lrp* RBS strength or the *eyfp* RBS strength was low, the fold-change was close to 1, meaning that these constructs hardly showed any sensor functionality. Six sensors with the lowest fold-change (below 1) showed a high fluorescence output both in presence and in absence of inducer. This result was unexpected. These sensors had a low *lrp* RBS strength in combination with a strong or medium *eyfp* RBS strength. Thus, results indicate that a low expression of *lrp* relieved the Lrp-based regulation of *eyfp*, resulting in expression both in presence and absence of inducer. This observation suggests that Lrp might function as both an activator and a repressor. Such a dual-regulatory function is known for the transcription factor AraC [28]. However, previous results showed that deletion of *lrp* reduced the export of L-isoleucine [29], so the exact regulatory mechanism should be investigated further.

**Fig. 6.**
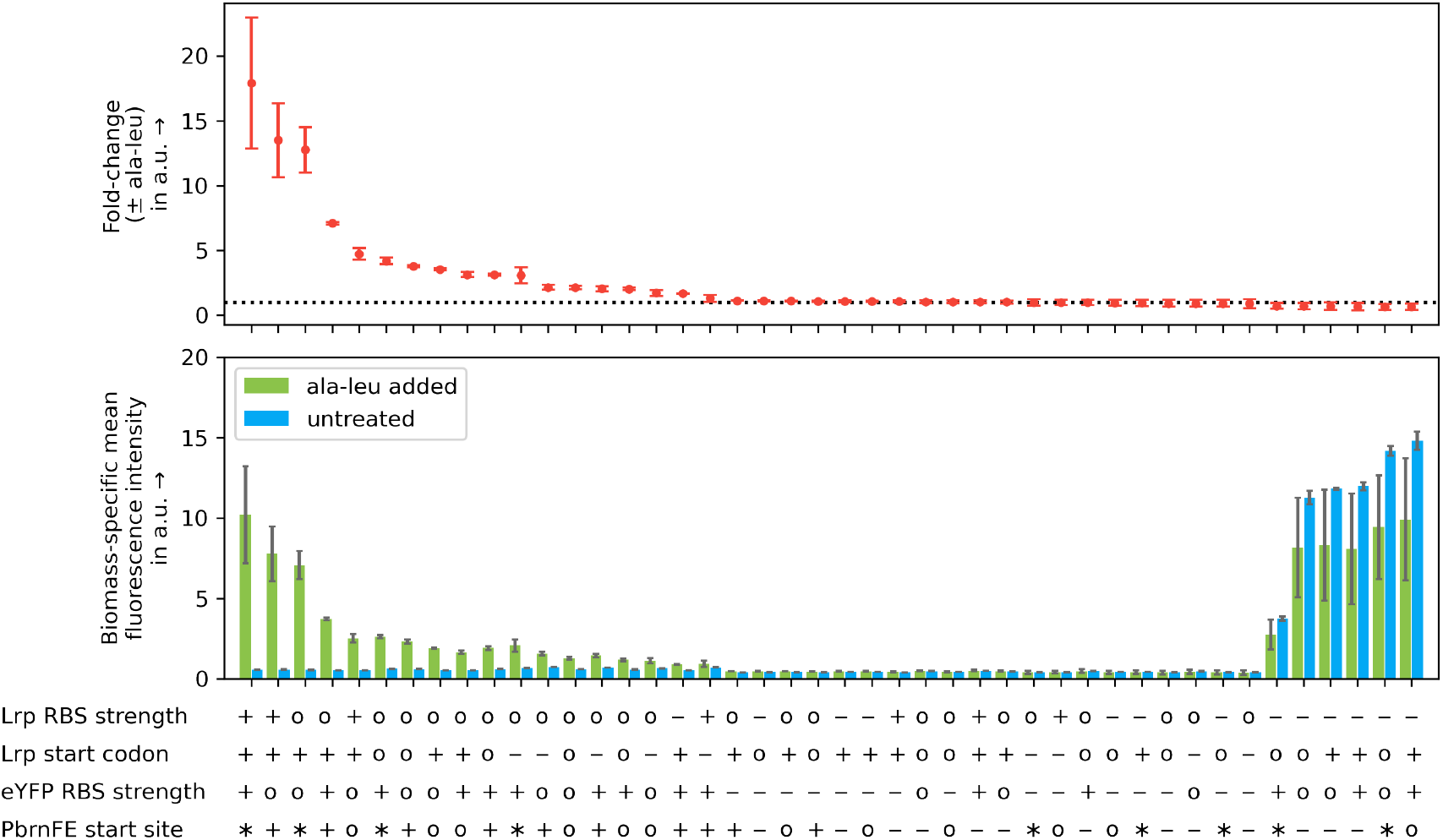
Screening results from the application example. Fold-change of each strain is calculated by dividing the biomass-specific fluorescence after 20 hours of growth with inducer (3 mM alanine-leucine dipeptide) by the biomass-specific fluorescence after 20 hours of growth without inducer. Dashed line indicates a fold-change of one. Higher values indicate a higher dynamic range of the sensor. Symbols used for strain characteristics: *lrp* RBS strength: −, weak | ◦, average (native) | +, strong; *lrp* start codon: −, weak | ◦, average | +, strong (native); *eyfp* RBS strength: −, weak | ◦, average (native) | +, strong; *brnF* promoter start site: −, −15 | ◦, 0 | +, +15 | ∗, +30. Mean and standard deviation of three biological replicates are shown.

### Considerations on the automated cloning workflow

The automated cloning workflow presented in this study still features some manual parts:

1. The preparation of competent *E. coli* cells, which is necessary for heat shock transformation. It has been shown that this procedure can be automated [30]. For the study presented here, this approach was not worthwhile since *E. coli* is only used as a shuttle system to carry the plasmid and perform conjugation into *C. glutamicum*. This task can be fulfilled by using a single *E. coli* strain, which can be manually prepared in bulk and stored by cryo-conservation.
2. The targeted *C. glutamicum* culture was prepared manually in this study. The reasoning here is the same as before. Since only one strain is used, it is actually more efficient to use a simple shake flask culture then setting up a robot for automated cultivation. Of course, this changes when multiple target strains are to be transformed, which conjugation easily allows to do, or when one aims for a completely autonomous platform. Then, an automated cultivation can be realized, *e.g.*, by using microbioreactors or microtiter plate-based cultivation [22].
3. The task of colony picking is done by hand. As discussed, dedicated devices are available for this operation [31]. They can be easily integrated into the existing workflow, further reducing the manual labor involved in molecular cloning.
4. The transfer of labware between the different devices used is done manually. Although not time-consuming, and automated transfer and orchestration of the different unit operations is necessary for a completely autonomous workflow. In this study, this challenge was partly alleviated by the long incubation times, shifting transfer of labware and start of operations into standard work hours.

Two different liquid handling systems were used for this publication: A low-cost device, the Opentrons OT-2, and a liquid handling platform based on a Tecan EVO200. To make the workflow as accessible as possible, it would be beneficial to only use the OT-2. For all the steps leading up to the conjugation, this can easily be realized by integrating the necessary modules provided by the manufacturer, *i.e.*, the temperature module, magnetic module, and thermocycler. Conjugation is more difficultt or realize, since it involves multiple steps using centrifugation. This cannot be done using the OT-2 which lacks a robotic manipulator arm, since labware must be moved in and out of a centrifuge. Simple sedimentation of *E. coli* and *C. glutamicum* did not yield any transformants (data not shown), presumably because the distance between donor and receptor organisms remained too large. A semi-automated procedure, where the OT-2 is responsible for liquid handling and the human operator for manual centrifugation, seems reasonable for this unit operation.

Regarding the efficiency of the presented workflow, two domains have to be discussed: The number of constructs successfully passing through each unit operation, and the time necessary to perform the operations. Most constructs were lost after the heat shock operation for creating transformed *E. coli* cells (Figure 7A). This could have multiple reasons, leaving room for further investigations: Either the heat shock operation itself had low efficiency, or the steps leading up to this operation did not work properly. Operations and reaction conditions were identical for all constructs except for the PCR reactions, where different primers were used to generate the biosensor variations. Thus, one can assume that this operation is responsible for the loss in constructs. This could possibly have been alleviated by introducing replicates, but since it is usually not possible to deal with all constructs in one pass of this workflow anyways, it would be most efficient to continue the workflow regardless of losses and queue the failed candidates for the next pass. This would work especially well when combined with scheduling approaches (see below).

**Fig. 7.**
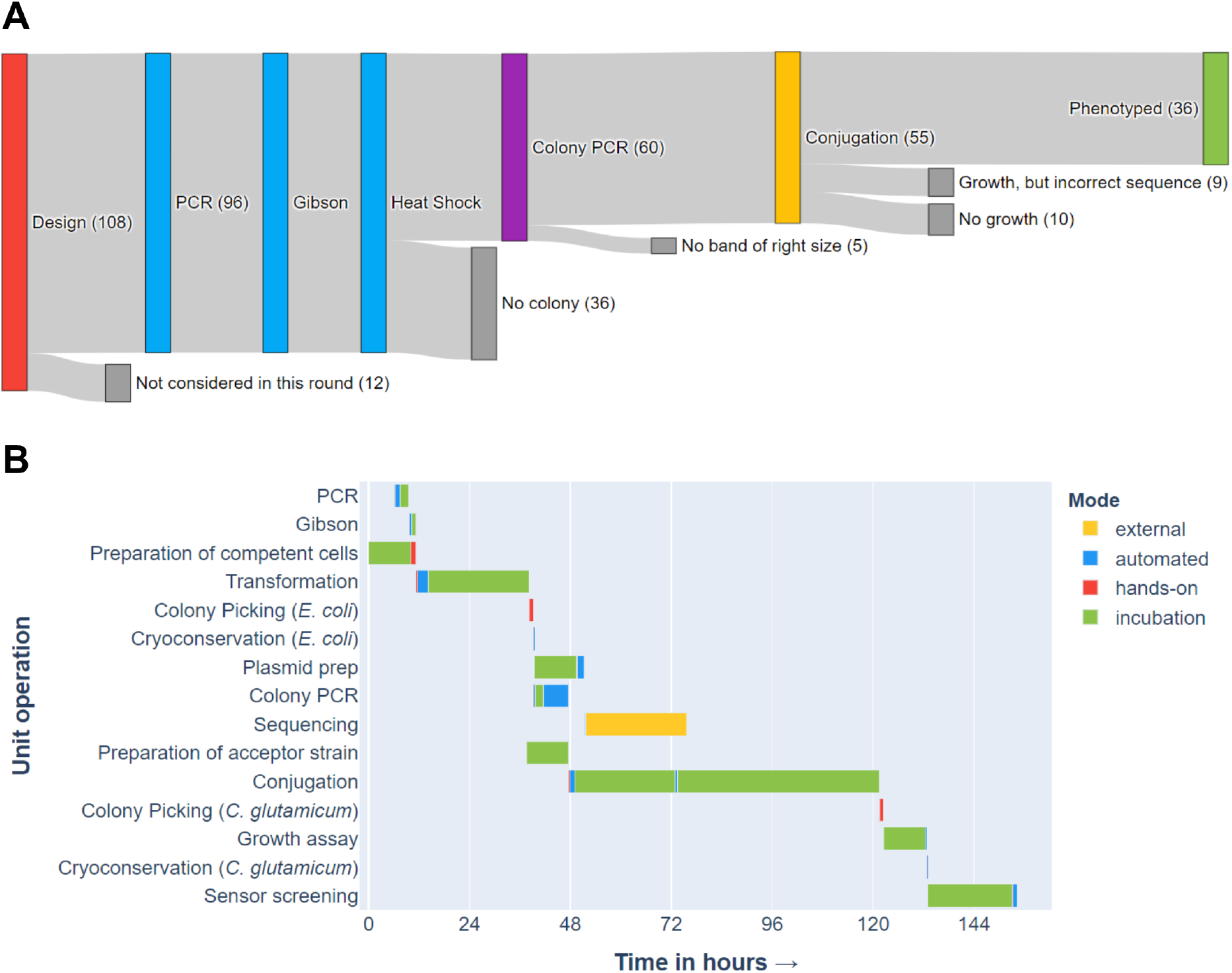
Efficiency of the automated cloning workflow. (A) Number of constructs passing each unit operation of the workflow. (B) Gantt chart of all unit operations.

Incubation times took most of the total workflow time (Figure 7B). This if mostly due to the relatively slow growth rate of the target organism, *C. glutamicum*, which is most prevalent in the conjugation step. But also *E. coli* operations are slowed down by the necessary incubation times. Switching to an even faster organism as a tool for molecular cloning, *e.g. Vibrio natrigens* [32], might be an option here, provided that a strain that is able to perform conjugation can be obtained. To increase throughput, especially on a fully autonomous platform, one could employ scheduling, preparing one microtiter-plate of constructs after the other and incubate them together. Since the whole workflow is designed for batches of 96 constructs, development of such a scheduled workflow is straightforward. The Gantt chart also demonstrates the reasoning behind the different feedback levels within the workflow: Colony PCR provides a quick first evaluation of the quality of the constructed *E. coli* strains, helping to use strains with a fully assembled plasmid for conjugation. Sequencing provides a detailed genotype but takes longer and is more costly. Therefore, this information is incorporated into the workflow at a later stage.

## Conclusions

The presented study shows the development of an automated workflow for high-throughput rational construction of plasmids and their subsequent transfer into an industrial-relevant target organism by conjugation. The workflow is described and characterized in-depth to be used as a basis for further developments. One novel unit operation, the automated conjugation of *E. coli* and *C. glutamicum*, enables high-throughput genetic engineering of this otherwise laborious to engineer microorganism. Most importantly, this process is not specific to one target species. It can rather be used for any microorganism susceptible to horizontal gene transfer via conjugation, with little to no effort for preparation of the target strain. Furthermore, recent work has shown the potential of conjugation to genetically engineer a wide range of established and novel hosts [33]. The workflow presented here is accompanied by a custom-made software tool to keep track of the constructed variants and their status. By applying the operations described in this study, an Lrp-biosensor library was constructed and characterized., resulting in the identification of a sensor with an improved dynamic range. Furthermore, the fact that several sensor variants showed output in the absence of effector hints at a reassessment of the function of Lrp as both an inducer and a repressor, thus providing novel fundamental insights. Building upon this workflow and filling the remaining gaps, an autonomous biofoundry for engineering of industrial microbial strains is within reach.

## Material and Methods

### Bacterial strains, plasmids, primers, and growth media

*E. coli* S17-1 [34] was used for plasmid propagation, storage and as donor strain for plasmid conjugation. Cultures were grown in Lysogeny Broth (LB) medium or on LB-agar plates at 37 °C. Supplementation with antibiotics was done, depending on the experiment, with 5 *µ*g mL^−1^ tetracycline (Sigma Aldrich, USA).

All *C. glutamicum strains* are derived from *C. glutamicum* ATCC 13032 [35] and were grown aerobically at 30 °C on Brain Heart infusion (BHI) (Difco Laboratories, Detroit, USA) agar plates (with 1.8 % (w v^−1^) agar). Supplementation with antibiotics was done, depending on the experiment, with 5 *µ*g mL^−1^ tetracycline (Sigma Aldrich, USA).

Defined CGXII medium [36] contained per liter of deionized water 20 g (NH_4_)_2_SO_4_, 1 g K_2_HPO_4_, 1 g KH_2_PO_4_, 5 g urea, 42 g MOPS, 13.25 mg CaCl_2_ · 2 H_2_O, 0.25 g MgSO_4_ · 7 H_2_O, 10 mg FeSO_4_ · 7 H_2_O, 10 mg MnSO_4_ · H_2_O, 0.02 mg NiCl_2_ · 6 H_2_O, 0.313 mg CuSO_4_ · 5 H_2_O, 1 mg ZnSO_4_ · 7 H_2_O, 0.2 mg biotin, 30 mg 3,4 dihydroxybenzoate (PCA), 20 g D-glucose and 5 mg tetracycline was used for *C. glutamicum* phenotyping experiments. Supplementation with 3 g L^−1^ alanine-leucine dipeptide (Bachem, Switzerland) was done depending on the experiment.

All constructed plasmids were derived from pEC-T18mob2 [27], an *E. coli*–*C. glutamicum* shuttle vector that contains the *E. coli* oriV, the *C. glutamicum* pGA1 ori and a tetra-cyline resistance marker. Plasmid pJC1-*lrp*-*brnF’*-*eyfp* [18] was used as template for all Lrp biosensor variant construction. For initial conjugation testing experiments, plasmids pEC-C18mob2 and pEC-S18mob2 were used [26]. Primers were ordered as custom DNA oligonucleotides from Eurofins Genomics (Ebersberg, Germany). All primers are listed in the Supporting Information (Table S1).

### Robotic platforms and 3D printing

In this study, two robotic platforms were used for automation of genetic engineering workflows: The Opentrons OT-2 (Opentrons Inc., New York, NY, USA) is a benchtop liquid handling device, capable of transferring liquids using electronic pipettes and disposable tips. To keep reagents and plates cooled on the robot’s deck, a rack which could be filled with ice and fitting adapters was designed and 3D printed (see Supporting Information Figure S1). Additionally, a BioShake 3000 elm (Quantifoil Instruments GmbH, Jena, Germany) was placed next to the system to provide quick access to shaking and heating capabilities by manually placing a microtiter plate on the device.

For more sophisticated processes, a customized Freedom EVO200 liquid handling system (Tecan, Männedorf, Switzerland) was used. The system provides the following capabilities: Liquid handling access to up to 16 plates and/or tube racks, cooling of up to 3 microtiter plates down to 4 °C, heating and shaking (BioShake 3000 elm, Quantifoil Instruments GmbH, Jena, Germany), centrifugation (4-5KRL, Sigma Laborzentrifugen GmbH, Osterode am Harz, Germany or Rotanda 460 Robotic, Andreas Hettich GmbH & Co. KG, Tuttlingen, Germany) and photometric measurements (Infinite M200, Tecan, Männedorf, Switzerland). Additionally, the liquid handling arm has access to a microtiter plate cultivation device (BioLector, m2p-labs GmbH, Germany).

3D printing was realized by fused-filament fabrication using an Original Prusa MK3S (Prusa Research a.s., Prague, Czech Republic) with polylactic-acid filament (DAS FILAMENT, Emskirchen, Germany). Design of 3D models was done using SolidWorks 2016 (Dassault Systèmes, Vélizy-Villacoublay, France).

### PCR

Automated polymerase chain reaction (PCR) was performed using the OT-2 liquid handling system and a qTOWER 2.2 (Analytic Jena, Jena, Germany) thermocycler. A microtiter plate holding the PCR reactions was prepared by distributing 15 *µ*L manually prepared PCR master-mix, containing Q5® High-Fidelity 2X Master Mix (New England Biolabs GmbH, Frankfurt am Main, Germany) and plasmid pJC1-*lrp*-*brnF’*-*eyfp* [18] as template, to each well and adding 1 *µ*L of each primer. All reagents were kept cool on ice during the pipetting process. The plate was then covered with foil (SILVERseal, Greiner Bio-One International GmbH, Kremsmünster, Austria) and placed into the thermocycler (qTOWER 2.2, Analytic Jena, Jena, Germany). Thermocycling was done according to manufacturer’s instructions.

Manual PCR was performed using a Biometra personal thermocycler (Analytic Jena, Jena, Germany) thermocycler and the Q5® High-Fidelity 2X Master Mix (New England Biolabs GmbH, Frankfurt am Main, Germany), according to manufacturer’s instructions.

### A. Gibson assembly

Gibson assembly [21] was done by combining 2 *µ*L of PCR product with 8 *µ*L of manually prepared Gibson assembly mastermix, containing 2 ng *µ*L^−1^ of NcoI-cleaved backbone (pEC-Tmob18-*lrp*-*eyfp*), in a microtiter plate using the OT-2 or EVO-200. All reagents were kept cool on ice or on the cooling carrier during the pipetting process. The plate was then covered with foil (SILVERseal, Greiner bio-one, Kremsmünster, Austria) and placed in a qTOWER 2.2 (Analytic Jena, Jena, Germany) thermocycler at 50 °C for 1 hour. Manual Gibson assembly of the pEC-Tmob18-*lrp*-*eyfp* backbone was done by combining 1 *µ*L of each PCR product (*lrp* and *eyfp*) with 3 *µ*L of EcoRI and BamHI cleaved backbone (pEC-Tmob18) and 5 *µ*L of Gibson assembly mastermix. All reagents were kept cool on ice and placed in a Biometra personal thermocycler (Analytic Jena, Jena, Germany) at 50 °C for 1 hour.

### Heat shock transformation of *E. coli*

*>E. coli* S17-1 [34] competent cells were prepared manually according to the rubidium chloride method [37]. Heat shock transformation was done using the EVO200 liquid handling system. The heating device was set to 42 °C and a V-Bottom plate (Greiner Bio-One International GmbH, Kremsmünster, Austria) was placed on it. 2 *µ*L per well of Gibson assembly product was distributed to a cooled (4 °C) microtiter plate. 25 *µ*L of competent *E. coli* S17-1 cells were added from a tube. The plate was incubated for 30 minutes. After incubation, the following procedure was done for each well of the microtiter plate, eight wells at a time: The complete volume of cell suspension was taken up by the liquid handler, pipetted into a well of the heated V-Bottom plate, was incubated for 30 seconds, was taken up again and transferred back to the cooled plate. After this procedure was done for the whole plate, 200 *µ*L BHI medium was added to every well. Afterwards, all wells were transferred to a 2 mL deep-well plate and 600 *µ*L BHI medium was added. This plate was shaken at 800 rpm and 37 °C for 60 minutes to provide cell recovery. For plating out, a spotting routine was employed: 12 round agar plates were placed on the robotic deck using a custom-made adapter (see Supporting Information). For each well, 2 spots with 5 *µ*L of the cell suspension were made. Afterwards, the deep-well plate was spun down using a centrifuge (4500 rpm, 5 minutes), supernatant was removed, and the cells were resuspended in 50 *µ*L LB medium. From this solution, another 4 spots per well, again with 5 *µ*L, were made.

### Incubation and colony picking

Agar plates were incubated for 16 h at 37 °C. For each construct, at most four colonies were picked by using a sterile toothpick, carefully touching the colony, dipping the tooth-pick into the master-mix for Colony PCR and intos LB medium for liquid culture.

### Plasmid preparation

For automated plasmid preparation, a ChargeSwitch NoSpin Plasmid Micro Kit (Thermo Fisher Scientific, Waltham, USA) was used on the EVO200 liquid handling system according to the manufacturer’s instructions. As magnet, a Magnetic-Ring Stand (Thermo Fisher Scientific, Waltham, USA) was used. Plasmid DNA was eluted using 50 *µ*L elution buffer. Quality was checked by analyzing a random sample (*n* = 8) using a Implen NanoPhotometer® P330 (Implen, Bayern, Germany). Sanger sequencing of plasmids was done by Eurofins Genomics (Ebersberg, Germany).

### Colony PCR, standard and capillary gel electrophoresis

Colony PCR thermocycling was performed using a qTOWER 2.2 (Analytic Jena, Jena, Germany) thermocycler and the OneTaq® 2X Master Mix with Standard Buffer (New England Biolabs GmbH, Frankfurt am Main, Germany), according to manufacturer’s instructions. PCR products were loaded on a 2 % (w v^−1^) agarose gel and gel electrophoresis was don for 40 minutes at 100 V. GeneRuler™ 100 bp Plus DNA-Ladder (Thermo Fisher Scientific, Waltham, USA) was used for quantification of PCR product sizes. Additionally, PCR products were analyzed by capillary gel electrophoresis using a MCE-202 MultiNA capillary gel electrophoresis device (Shimadzu Corp., Nakagyo-ku, Kyōto, Japan) according to manufacturer’s instructions.

### Conjugation of *E. coli* and *C. glutamicum*

Conjugation was performed by preparing liquid cultivations of *E. coli* harboring the assembled plasmids and *C. glutamicum*. *E. coli* was cultivated using a deep well plate filled with 200 *µ*L LB medium per well, shaken at 900 rpm, 37°C, overnight. Prior to conjugation, 800 *µ*L LB medium was added per well (1 : 5 dilution), and the plate was shaken at 900 rpm, 37 °C, for 3 hours. For *C. glutamicum* a preculture was prepared by using a 500 mL shake flask filled with 50 mL BHI medium, shaken at 250 rpm, 30 °C, overnight. Prior to conjugation, this culture was transferred to a 1000 mL shake flask filled with 50 mL fresh BHI medium (1 : 2 dilution). This culture was shaken at 250 rpm, 30 °C, for 3 hours. Right before starting the conjugation procedure, the *C. glutamicum* culture was placed in a 48.5 °C water bath for 9 minutes, to inhibit the restriction modification system [38]. Both cultures were placed on the robotic deck of the EVO200 system. For *C. glutamicum*, a through was used. From each *E. coli* culture, 250 *µ*L were transferred to 1 mL deep well plate, and 750 *µ*L of *C. glutamicum* culture were added. The plate was then centrifuged (4500 rpm, 5 minutes), the supernatant removed, and the pellet resuspended in LB media by pipetting up and down 10 times with 950 *µ*L volume. The plate was then again centrifuged (4500 rpm, 5 minutes) and the supernatant partly removed, so that 300 *µ*L were left on top of the pellet. The plate was covered with gas-permeable sealing foil (m2p-labs, Baesweiler, Germany) and placed in an incubator for 20 hours. After incubation, each well was thoroughly mixed by pipetting up and down 10 times with 950 *µ*L BHI. From each well, two spots with 5 *µ*L were plated out on an BHI agar plate containing antibiotics tetracylin (5 *µ*g mL^−1^) and nalidixin (50 *µ*g mL^−1^). The plate was centrifuged (4500 rpm, 5 minutes), the supernatant removed and each well resuspended with 50 *µ*L BHI medium. From this concentrated suspension, four additional spots with 5 *µ*L were prepared on agar plates. After approximately 40 hours of incubations, single *C. glutamicum* clones were picked into BHI medium containing antibiotics tetracylin (5 *µ*g mL^−1^) and nalidixin (50 *µ*g mL^−1^) for storage and phenotyping.

### Phenotyping of *C. glutamicum* strain variants

Cultivations for phenotyping of *C. glutamicum* strain variants were performed using specialized 48-well microtiter plates (FlowerPlate, m2p-labs GmbH, Baesweiler, Germany). The EVO200 system was used to distribute defined CGXII medium with or without supplementation of 3 mM alanine-leucine dipeptide to the plates. Inoculation using 10 *µ*L of thawn cryocultures per well was also done using this robot. Plates were then covered with gas-permeable sealing foil (m2p-labs GmbH, Baesweiler, Germany), transferred to an incubator (TiMix / TH 15, Edmund Bühler GmbH, Bodelshausen, Germany) set to 30 °C and shaken at 1400 rpm.

After 20 hours of incubation, a sample of 250 *µ*L was drawn from each well and transferred to a multititer plate (PS, F-Bottom, transparent, Greiner Bio-One International GmbH, Kremsmünster, Austria) for subsequent measurement of absorption at 600 nm and fluorescence ( excitation 488 nm, emission 525 nm).

### Computational methods and TACO

The whole workflow was accompanied by data management using a tailor-made toolbox named TACO (Tool for Automated Cloning). TACO is written in Python 3.8 and available to the general public under MIT license (see Availability). TACO uses pandas [39] and numpy [40] for most of its functionality. User input is done directly via Python or using Microsoft Excel (Microsoft Corp., Seattle, WA, USA). Graphviz [41] is used for visualization within TACO. Matplotlib [42], Seaborn [43] and plotly (Plotly Inc., Montreal, Quebec, Canada) were used for additional plotting.

## Availability

The source code for the TACO package and data from this study as an application example is freely available at https://github.com/MicroPhen/taco. Additional data can be made available on request.

## ACKNOWLEDGEMENTS

Funding was received by the German Federal Ministry of Education and Research (BMBF, projects: “BioökonomieREVIER_INNO: Entwicklung der Modellregion BioökonomieREVIER Rheinland”, grant no. 031B0918A) and by the Helmholtz Association (grant W2/W3-096).

## Supporting Information

**Table S1.**
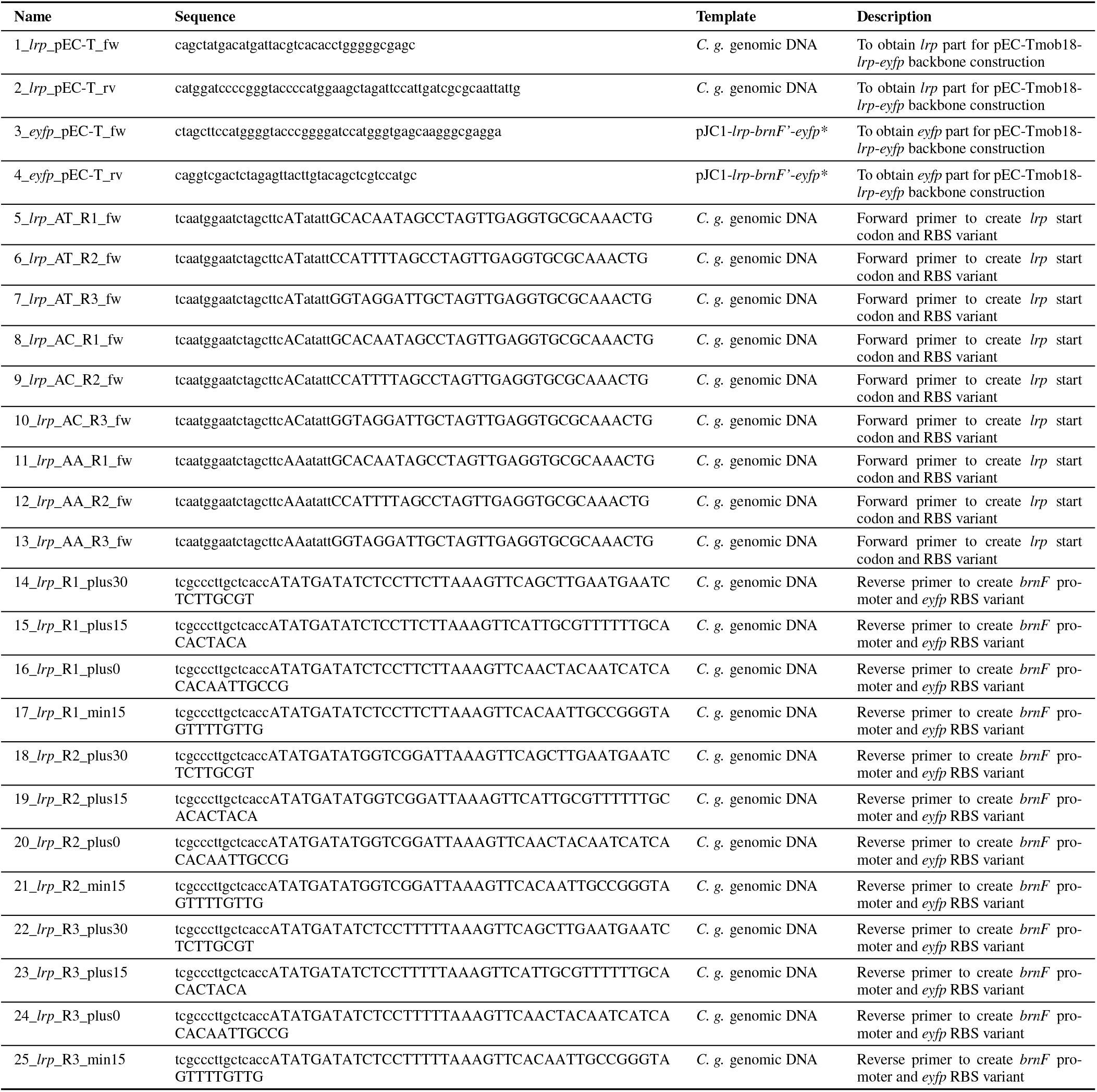
Primers used in this study.

**Fig. S1.**
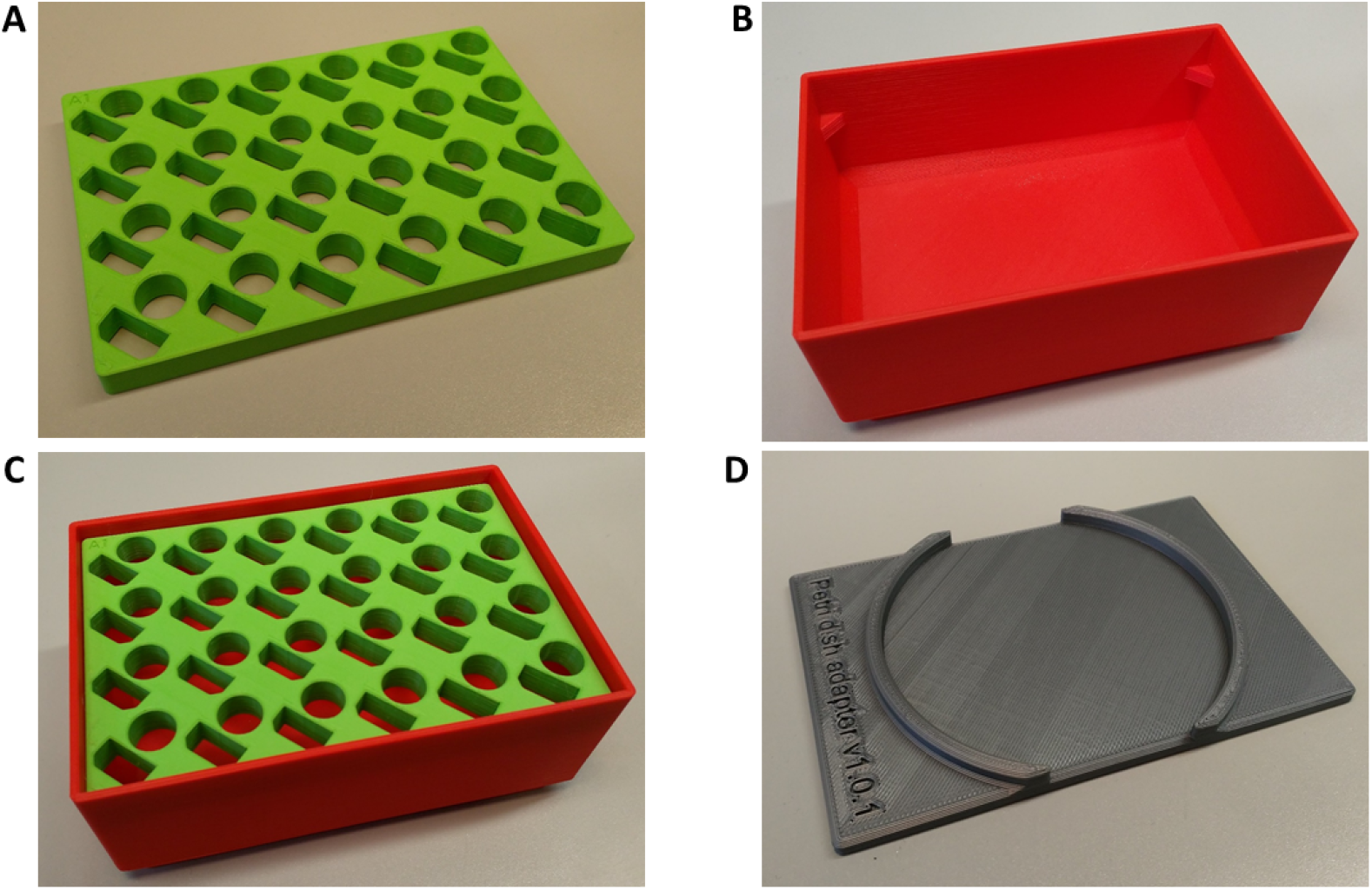
Images of 3D printed parts used in this study. All files are freely available at https://github.com/MicroPhen/taco. (A) Plate for placement of 1.5 mL or 2 mL reaction tubes. Filename: “reaction-tube-adapter-for-ot-cc.stl”. (B) Trough for crushed ice with SBS format base and SBS format opening. Filename: “ot-cooling-carrier.stl”. (C) Assembly of reaction tube holder A in crushed ice trough B. (D) Petri dish adapter for 100 mm Petri dishes. Filename: “petri_dish_adapter.stl”.

